# Time-frequency embedding with contrastive pre-training allows sub-second seizure detection

**DOI:** 10.64898/2026.01.21.700017

**Authors:** Helena A. Merker, Isabella Dalla Betta, Matthew A. Wilson, Francisco J. Flores, Emery N. Brown

## Abstract

Rapid and accurate detection of electrographic seizures is critical for both clinical diagnosis and neuroscience research. Although seizure identification is commonly performed in the time domain, analysis in the time-frequency domain provides a more comprehensive representation of seizure characteristics. In this study, we present a 3D convolutional neural network (CNN) that incorporates a trainable continuous wavelet transform (CWT) layer, enabling adaptive time-frequency feature learning directly from raw EEG. To address common data challenges, we augment the 3D CNN for pre-training with contrastive learning, comparing contrastive predictive coding (CPC) against bidirectional contrastive learning (BiCL). On single-channel and multi-channel data, the standard 3D CNN outperformed both a 2D CNN with pre-computed CWT and a 1D CNN that processes raw signals, achieving *>*95% accuracy down to 0.5-second segments. Compared to the standard 3D CNN, the 3D CNN with BiCL pre-training showed superior performance in both low-data and class imbalance scenarios. Further experiments involving band-pass filtering and temporal shuffling revealed that classification is driven primarily by low-frequency patterns and statistical features rather than temporal dependencies. The proposed framework also maintained *>*90% accuracy with moderate noise and downsampling applied to inputs, as well as when cross-subject generalization was evaluated using held-out subjects. We show that a 3D CNN with a trainable CWT layer and BiCL pre-training enables accurate sub-second seizure detection and effectively mitigates data limitations common in clinical settings. This work demonstrates that time-frequency embedding within CNNs, augmented by self-supervised pre-training, offers a promising path toward architectures for sub-second seizure detection in the presence of practical limitations of real-world scenarios.

## Introduction

Fast identification of seizures in the brain has been a desirable goal in both research and clinical practice since the early days of electroencephalography (EEG).^1^ In clinical settings, efficient and accurate diagnosis of electrographic seizures is essential.^2, 3^ Rapid EEG assessments can significantly reduce hospitalization duration and mitigate the risks associated with missed or misidentified seizures.^4, 5^ In research settings, quick seizure identification facilitates the accurate study of the neural mechanisms that underlie seizures and allows testing of strategies to promptly halt seizures in animal models, whether in open- or closed-loop fashion.^6^ Most seizure detection algorithms rely upon large training datasets; however, the manual labeling of EEG data to generate these datasets is costly and labor-intensive.^7, 8^ Another challenge is that the stochastic nature of EEG signals and the prevalence of artifacts can lead to low inter-rater reliability.^9^ Moreover, machine learning algorithms commonly rely on the training and testing data sharing the same distribution, an assumption that typically does not hold in generic applications of EEG signal recognition.^10^ In recent years, deep learning techniques have revolutionized EEG analysis by automating feature extraction.^11^

Convolutional neural networks (CNNs) have been successfully applied to raw EEG signals, learning representations that capture spatial dependencies.^12^ However, approaches that directly process raw signals can be challenged by noise and artifacts.^13, 14^ Researchers have increasingly turned to time-frequency representations as pre-processing steps,^15–17^ as it supplies the network with the explicit temporal and spectral content of the signal.^18^ Among the various time-frequency analysis techniques, the continuous wavelet transform (CWT) has been proven effective for EEG analysis.^19^ The CWT decomposes signals over a continuum of scales, offering a high-resolution, multiscale view that can capture both high-frequency bursts and low-frequency oscillatory components—features that are critically important for identifying seizures.^20, 21^ Various approaches use the scalograms from the CWT as input to 2D CNNs.^22–25^

When there is significant data scarcity or class imbalance, self-supervised pre-training with contrastive predictive coding (CPC) can effectively capture meaningful features from unlabeled data.^26^ Since CPC simply requires that observations be ordered, for instance along temporal dimensions, the technique has been applied to a variety of modalities including images, speech, and natural language.^27^ CPC representations have been shown to enhance the performance of downstream tasks such as emotion recognition and speech classification,^28, 29^ and in the image domain, CPC has been shown to improve classification accuracy when there are small amounts of labeled data.^27^ The CPC framework includes an encoder network that maps input sequences to latent representations, which are then processed by an autoregressive model, such as a transformer with masked self-attention, that can only access past information when predicting future states.^26, 30^ While CPC relies on autoregressive architectures to encode context, other approaches have used bidirectional transformers on contrastive learning objectives.^31–33^

In this work, we leverage the strengths of wavelet-based time-frequency analysis and deep learning to achieve rapid and accurate seizure identification with sub-second precision. We test our approach on a dataset of seizures in mouse EEG that we collected in a previous work.^34^ We propose a 3D CNN that integrates the wavelet transform directly into the architecture, enabling custom filters trained for the task of seizure detection. To further improve the generalizability of the model, we also apply pre-training with contrastive learning, replacing the autoregressive model within the CPC framework with a bidirectional transformer. Our results indicate that, even when only a small amount of labeled data is available for fine-tuning, seizures can be reliably identified. This work represents a step forward towards solving the general challenge of developing high-performance seizure detection systems for diverse clinical and research environments.

## Methods

### Data Collection

All experiments were performed in accordance with MIT IACUC (protocol # 2303000480) and National Institutes of Health guidelines. C57BL/6J mice (n = 10, 5 male) were implanted with bilateral bipolar tungsten electrodes for stimulation (E363T/3-2TW/SPC, P1 Technologies) aimed at the central thalamus and stainless steel screws for EEG recording over the frontal and parietal cortices (8209, Pinnacle Technology). Constant current stimulation (20–200 µA; STG-8000, Multichannel Systems) consisted of biphasic, cathodal first, charge-balanced, 100-µs square pulses separated by a 100-µs iso-electric period. The EEG was recorded at 2713 Hz, band-pass filtered (0.1–500 Hz; Neuralynx), and reviewed independently by two researchers to identify electrographic seizure activity present in at least one electrode. For a detailed description of these methods, see Flores et al.^34^ Central thalamic stimulation induced electrographic seizures during 53 recording sessions for a total of 82 seizures, lasting from 0.5 to 55 s. 74 seizures (total duration: 1495 s) included both frontal and parietal EEG channels, but due to signal quality issues, 7 (189 s) included only the frontal channel and 1 (28 s) included only the parietal channel. Normal, wake EEG activity was extracted from the beginning of each recording session before any seizure activity occurred, yielding 50 non-seizure EEGs (1255 s) with both frontal and parietal channels, 2 (89 s) with only frontal, and 1 (28 s) with only parietal. The EEG traces and corresponding scalogram of a representative 22-s seizure are shown in Fig **1**A. To expand the parietal dataset to include sleep data, 44 sleep EEGs (1245 s) were acquired in additional recording sessions and only periods of non-REM sleep were used.

**Figure 1:**
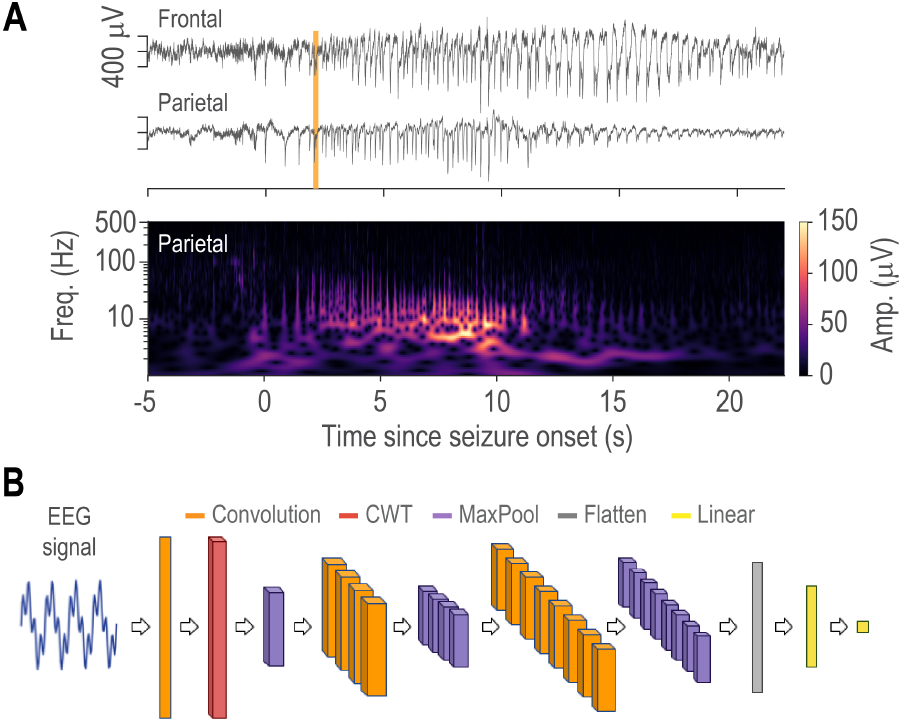
Example seizure and 3D CNN model architecture. (A) The top two traces show parietal and frontal EEG corresponding to a 22-s seizure low-pass filtered at 500 Hz. Seizure onset occurs at t = 0 (orange line in top panel). The vertical scale (400 *µ*V) is the same for both traces. The lower panel shows the CWT scalogram of the seizure trace on top. (B) Architecture of the 3D CNN. A 1D convolutional layer (orange) handles initial feature extraction, followed by a CWT layer (red) that captures complex-valued wavelet features across multiple scales. Subsequent 3D convolutional (orange) and max-pooling layers (purple) are followed by a flattening (gray) and linear layers (yellow), refining features and enabling classification. Each 3D convolutional layer, as well as the first linear layer, are followed by ELU activation.

### CNN Architecture

Seizure detection on time-domain EEG signals requires modeling the signals to extract meaningful features.^8^ Our architecture integrates a trainable CWT layer within a 3D CNN (Fig **1**B). We incorporate the CWT into our model to perform feature extraction by multi-scale analysis of the input signals and let its parameters adapt during the training process, allowing the model to learn the optimal parameters for the task of seizure detection. We compare the 3D CNN to two other architectures to assess the impact of different inputs. The first comparison model is a 2D CNN that takes as its input the CWT computed outside the network. When compared to the 3D CNN, the 2D CNN examines whether incorporating the CWT as a layer or as a pre-processing step yields better results. The second comparison model is a 1D CNN that takes as its input the EEG time series without any time-frequency decomposition. When evaluated against the other models, this approach determines whether adding a time-frequency decomposition, trainable or not, improves performance.

The first layer within the 3D CNN is a 1D convolutional layer with a kernel size of 5 and a stride of 3, which reduces the temporal dimension and extracts the initial features from the raw EEG signal. This output is subsequently passed to the CWT layer, after which a 3D max-pooling layer downsamples the time-frequency feature space. Two subsequent sets of 3D convolutional and max-pooling layers are then applied. The first set consists of 4 filters with a kernel size of (4, 8, 1) and is scaled via ELU activation.^35^ The second set increases the number of filters to 8, maintaining the same kernel size and activation function as the first set. All max-pooling layers use a kernel size of (2, 4, 1). Dropout layers with probabilities of 0.25 and 0.5 are added after each set of layers to mitigate overfitting. The flattened output is then passed through two fully connected layers. The first dense layer reduces the dimension to 140 and uses ELU activation. The second dense layer has a single output, after which sigmoid activation is applied to determine the likelihood of seizure occurrence.

The 1D CNN begins with a 1D convolutional layer containing a single filter with a kernel size of 5 and a stride of 3, followed by ELU activation and max-pooling with a kernel size of 4. The network then applies two additional sets of convolutional layers with ELU activations and max-pooling. The first set uses 4 filters with a kernel size of 6, while the second set increases to 8 filters with the same kernel size. Similarly, the 2D CNN begins with a 2D convolutional layer using a single filter with a kernel size of (1, 5) and a stride of (1, 3), followed by ELU activation and max-pooling with a kernel size of (2, 4). Subsequently, two sets of convolutional layers are applied, each of which is followed by ELU activation and max-pooling. The first set employs 4 filters with kernel size of (4, 8), while the second set uses 8 filters with the same kernel size. In both the 1D and 2D CNNs, dropout layers with probabilities of 0.25 and 0.5 are incorporated after each set of layers to avoid overfitting. Both networks also conclude with two fully connected layers: the first reduces the dimensions to 140 with ELU activation and the second produces a single output for binary classification.

### The Continuous Wavelet Transform Layer

The CWT is a convolution between the input signal and a mother wavelet. A matrix of coefficients is obtained by sweeping across different scales and translations of the mother wavelet, where local maxima, aligned across scales, form ridges that highlight peak properties.^36^ Unlike the Fourier and Hilbert transforms, the CWT offers variable time-frequency resolution, making it especially effective for analyzing non-stationary and transient signals.^37^ The use of multiple scales captures signal characteristics across different frequency bands; high-frequency wavelets capture fine, detailed features in the signal, while low-frequency wavelets display large, coarse trends.^38^ For the signal *x*(*t*), the equation of the CWT *W*_*x*_(*a, b*) at scale *a* and time location *b* is given in Eq 1, where Ψ(*t*) is the mother wavelet.^39^ We chose a complex Morlet wavelet as the mother wavelet, which is defined as the product of a complex sine wave and a Gaussian window where *f* is the frequency and *σ* controls the width of the Gaussian (Eq 2).^40^

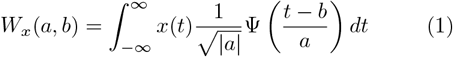

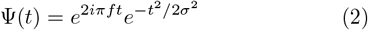

We set the frequency *f*, sigma *σ*, and filter length *l*, one for each scale, as trainable parameters. The real and imaginary parts of the wavelet are computed for every scale, padded to the same length, and subsequently stacked into two tensors. Each tensor is of the shape (*s*, 1, *l*_*max*_), where *s* is the number of scales and *l*_*max*_ is the maximum filter length. The input tensor to the CWT module is of the shape (*c, t*), where *c* is the number of channels and *t* is the number of time points. To process each channel as a separate signal with one channel, the input signal is reshaped to (*c*, 1, *t*) and then convolved with the wavelet filters along the time axis. The resulting real and imaginary components are each of the shape (*c, s, t*) and are stacked along a new axis to form an output tensor with dimensions (*c, s, t*, 2).

### Contrastive Learning

The theory of predictive coding proposes that the brain uses a generative model of the environment to make predictions of the future, with the goal of minimizing the error between these predictions and the true sensory inputs.^41^ CPC mimics this process by predicting future latent states and updating the model by comparing the predictions to the actual future latent states.^42^ The over-arching aim of CPC is to learn abstract, global representations of signals rather than low-level, high-dimensional ones.^43^ To this end, we augment the 3D CNN for pretraining with contrastive learning, a fully self-supervised approach that learns representations directly from the data without requiring any manual labels. This work compares two pre-training strategies: first, a CPC approach employing an autoregressive transformer backbone with future-position masking to ensure predictions rely solely on past context; second, a bidirectional contrastive learning (BiCL) approach without masking to investigate whether fully contextualized representations produce superior embeddings. By evaluating both causal and non-causal variants, we aim to determine whether directional constraints improve representation robustness when labeled seizure data for fine-tuning is limited or skewed towards the non-seizure class.

For contrastive pre-training, the input signal is first processed through the 3D CNN described in Fig **1**B, stopping after the last max-pooling layer. This time-frequency output is reshaped into a sequence of *T* latent vectors *z*_*t*_ ∈ ℝ^16^. Each latent vector *z*_*t*_ is linearly projected into a hidden dimension *d* = 32, after which a sinusoidal positional encoding is added elementwise as defined in Eq 3-4, where *p* is the position and *i* is the dimension.^44^ A transformer, which comprises three layers with four attention heads each, produces context vectors *c*_*t*_ ∈ ℝ^32^. In the CPC variant, an upper-triangular attention mask enforces autoregressive prediction; in the BiCL variant, no mask is applied, allowing each position to attend to both past and future context. Each original latent *z*_*t*_ is processed through a learned sigmoid gate—comprising a single linear layer followed by sigmoid activation—whose output is element-wise multiplied with *z*_*t*_ to yield a calibrated latent 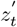. This gating mechanism allows the model to learn feature-wise scaling factors between 0 and 1. Next, the sequences of calibrated latents and transformer context vectors are each passed through parallel two-layer projection MLPs: the first layer maps the input (16 for latents, 32 for contexts) to a hidden dimension of 32, followed by layer normalization, Gaussian Error Linear Unit (GELU) activation, and dropout with probability of 0.3; the second layer then projects down to a 16-dimensional output, with a final layer normalization. We denote the MLP outputs as *u*_*t*_ ∈ R^16^ for the projected calibrated latents and *v*_*t*_ ∈ R^16^ for the projected context vectors.

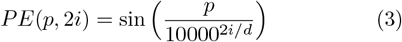

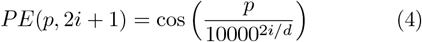

We optimize the model using InfoNCE loss, a contrastive objective that is based on Noise-Contrastive Estimation.^26^ InfoNCE optimizes the mutual information between a context vector and its corresponding future embedding by learning representations where positive pairs have high similarity relative to negative pairs. For each batch of *N* signals, at each time step *t* and each prediction step *k*, we treat (*v*_*i,t*_, *u*_*i,t*+*k*_) as the positive pair for sample *i*, and {(*v*_*i,t*_, *u*_*j,t*+*k*_)}_*j*≠*i*_ as the negatives. Denoting the temperature parameter *τ*, the loss for sample *i*, time *t*, and step *k* is given by Eq 5. The total loss is computed by averaging over all samples *i* ∈ {1, …, *N*}, all valid time steps *t* ∈ {0, …, *T* − *k* − 1}, and all prediction steps *k* ∈ {1, …, *K*}, where *K* = 10 is the maximum prediction horizon.

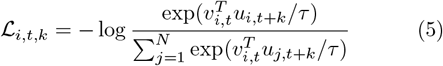

For pre-training, we utilized the CHB-MIT dataset, a comprehensive collection of scalp EEG recordings sampled at 256 Hz from pediatric patients with intractable seizures.^45, 46^ We extracted two electrode-pair channels: F7–T7 (frontal), which is the voltage difference between the left frontal electrode F7 and the left temporal electrode T7, and T7–P7 (parietal), which is the difference between T7 and the left parietal electrode P7.

## Results

To investigate the contribution of different cortical regions to seizure detection, we conducted experiments using data from the frontal cortex, parietal cortex, and both regions together. Each configuration was analyzed using the three CNN architectures (1D, 2D, and 3D), with the dual-cortex models modified to accommodate two-channel inputs. We divided each EEG into non-overlapping 2.0-second intervals, discarding the remaining data at the end of each recording. We also excluded recordings that were too brief (*<* 2 s) to generate any segments. This pre-processing resulted in frontal, parietal, and dual-cortex datasets of sizes 1450, 1345, and 1317 segments, respectively. The CWT frequency range of the 3D CNN was initialized as the linear space between 0.1 and 500 to match the EEG input. For all scales, sigma was set to the initial hyperparameter value of 5 while the filter length was initialized to 64 time points. In this work, all experiments use 8 scales. For the 2D CNN, the CWT was computed for each EEG segment, yielding complex coefficients, from which the magnitudes were extracted. Across the frontal, parietal, and dual-cortex datasets, the standard 3D CNN achieved higher accuracies than all other CNNs, suggesting that incorporating a CWT layer is the most effective approach for feature extraction (Fig **2**A). Comparing the different regions, the models trained on the frontal cortex data showed lower accuracies: (3D: 92.7 ± 2.5%; 2D: 87.9 ± 3.8%; 1D: 88.8 ± 4.2%) than those trained on the parietal cortex data (3D: 96.9 ± 1.4%; 2D: 92.9 ± 2.9%; 1D: 92.9 ± 3.9%). Moreover, the inclusion of both cortices had similar results to those of the parietal models (3D: 97.3 ± 1.4%; 2D: 93.0 ± 3.6%; 1D: 94.9 ± 2.1%), with the largest increase observed in the 1D CNN. These metrics indicate that, compared to the frontal cortex, the parietal cortex contains more informative features for seizure detection. We further investigated the error rates of each of the parietal CNNs. The 3D CNN showed the lowest percentage of false negatives at 2.8%, compared to 6.2% and 6.7% for the 2D and 1D CNNs, respectively (Fig **2**B).

**Figure 2:**
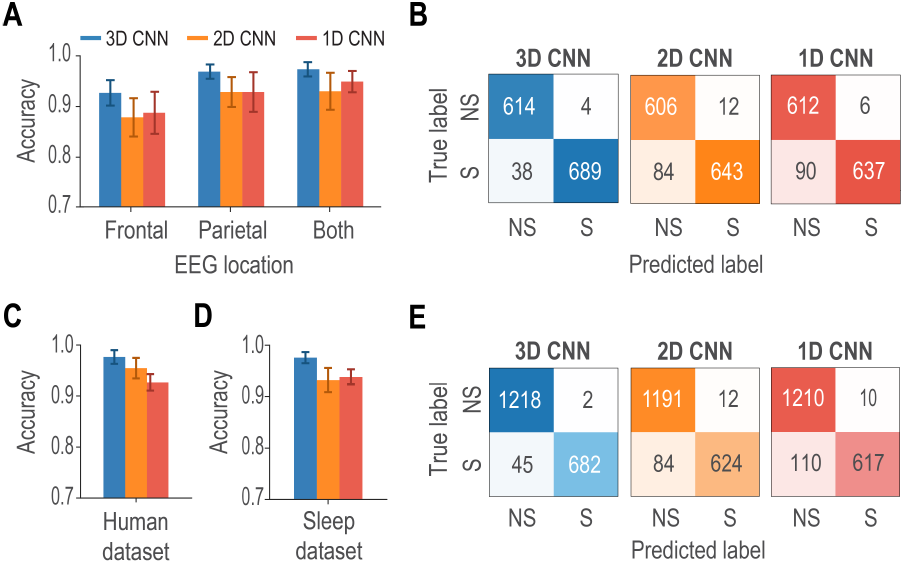
Performance of the models with different cortical data. (A) Mean accuracy of the 3D CNN (blue), 2D CNN (orange), and 1D CNN (red) architectures trained on frontal cortex data, parietal cortex data, or data from both cortices, evaluated through 10-fold cross-validation. Error bars correspond to the standard deviation of the mean. (B) Confusion matrices of the CNN architectures (same colors as in A) trained on parietal cortex data. NS and S denote non-seizure and seizure classes, respectively. (C) Mean accuracy of the 3D, 2D, and 1D CNN architectures trained on the Bonn dataset, evaluated through 10-fold cross-validation. (D) Mean accuracy of the 3D, 2D, and 1D CNN architectures trained on the expanded parietal cortex dataset (with sleep data added as additional non-seizure segments), evaluated through 10-fold cross-validation. (E) Confusion matrices of the CNN architectures trained on the expanded parietal cortex dataset.

Seizure and sleep EEG share overlapping spectral and temporal characteristics, making sleep an important physiological state for testing the efficacy of seizure detection. We next examined whether our approach could distinguish seizures from sleep activity. Each parietal sleep EEG was divided into non-overlapping 2.0-second segments, adding 602 non-seizure segments to the parietal cortex dataset. When trained and tested on this expanded dataset, the 3D CNN continued to outperform the 1D and 2D CNNs (Fig **2**D). However, all models showed an increase in seizures being misclassified as non-seizures, even though the total number of seizures remained constant (Fig **2**E). With the inclusion of sleep data, the dataset included a broader range of non-seizures that overlapped spectrally with seizures, resulting in the models becoming more conservative in assigning the seizure label. We additionally evaluated whether the 3D CNN remained advantageous across species by training and testing the CNNs on the Bonn dataset, a widely-used human EEG benchmark.^47^ Each sample was divided into non-overlapping 2.0-second segments at the sampling rate of 173.61 Hz and zero-padded to 5426 time points to maintain consistent input dimensions. The 3D CNN again achieved the highest accuracy, demonstrating that the architectural advantages generalize to humans despite cross-species differences (Fig **2**C). For the remaining results in this work, we used the original mice dataset without the inclusion of sleep EEGs.

In both low data and class-imbalanced scenarios, self-supervised pre-training provides a robust foundation for learning generalizable representations. In this work, we compare two self-supervised pre-training approaches: CPC, which enforces autoregressive context via masking in the transformer, and BiCL, which allows bidirectional context without any masking. For pre-training, recordings from the CHB-MIT dataset^46^ were segmented into contiguous, non-overlapping windows of 5426 time points (approximately 21.2 s), yielding 165917 total segments. Each model was pre-trained on CHB-MIT data acquired from the cortices corresponding to the fine-tuning data. We evaluated model performance under two conditions: low-data and class imbalance. In the low-data scenario, models were fine-tuned on a random 10% subset of our data, with the remaining data reserved for testing. For the class imbalance scenario, we simulated realistic clinical conditions where seizure events are rare by fine-tuning on a random 90% of non-seizure data but only a random 10% of seizure data, with the remaining data allocated to testing. Under low-data conditions, BiCL pretraining demonstrated an advantage for single-channel models versus CPC on frontal cortex data (BiCL: 92.2 ± 1.8%; CPC: 86.0 ± 5.4%) and on parietal cortex data (BiCL: 93.8 ± 1.2%; CPC: 90.5 ± 3.2%) (Fig **3**A). However, when both cortices were utilized, the performance gap between pre-training strategies diminishes. Similarly, under class imbalance conditions, both pre-training approaches yielded comparable performance, suggesting that the benefits of bidirectional context are most pronounced in single-channel, low-data scenarios.

**Figure 3:**
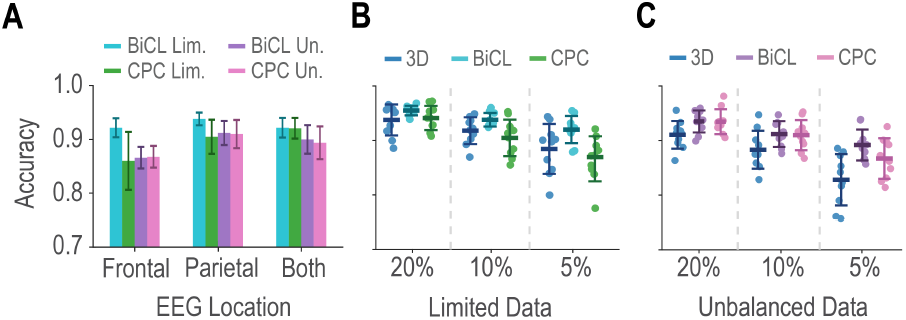
Performance of the fine-tuned models in low and unbalanced data conditions. (A) Mean accuracy of the 3D CNN architectures on frontal cortex data, parietal cortex data, or data from both cortices, across 10 trials with BiCL and CPC pre-training and two fine-tuning conditions: low-data (10% of available data, light blue for BiCL and green for CPC) and class imbalance (90% of non-seizure, 10% of seizure data, purple for BiCl and pink for CPC). (B) Performance of parietal 3D CNN models across varying levels of low data. Models were fine-tuned using 20%, 10%, and 5% of available training data, comparing baseline (no pre-training, dark blue), BiCL pre-training (light blue), and CPC pre-training (green). (C) Performance of parietal 3D CNN models under class imbalance conditions. Models were fine-tuned with 90% of non-seizure data and varying percentages of seizure data (20%, 10%, and 5%), comparing 3D CNN (blue), BiCL pretraining (purple), and CPC pre-training (pink). The vertical axis is the same in all plots.

Due to the higher accuracies of the CNNs trained on the parietal cortex data compared to those trained on the frontal cortex data, all subsequent experiments were performed using the parietal cortex data. Under fine-tuning with random 20%, 10%, and 5% subsets of the data, the 3D CNN with BiCL resulted in mean accuracies of 96%, 94%, and 92% (Fig **3**B). In comparison, the standard 3D CNN baseline resulted in lower accuracies of 94%, 92%, and 88%. No general performance improvement was observed when comparing pre-training with CPC to the baseline. Additionally, we investigated the parietal cortex CNNs under unbalanced training datasets that consisted of 90% of non-seizure data and 20%, 10%, and 5% of seizure data, with the remaining data reserved for testing. In contrast to the low-data experiments, both methods of pre-training resulted in higher accuracies than those of the baseline (Fig **3**C).

While CNNs are effective at identifying intricate patterns, interpreting the underlying features that they rely on remains a significant challenge. To identify the minimum temporal resolution necessary for reliable classification, we evaluated CNN performance across progressively shorter EEG segments. We segmented parietal cortex recordings into durations ranging from 2.0 s to 0.01 s (2.0, 1.5, 1.0, 0.5, 0.1, 0.05, and 0.01 s), excluding recordings shorter than 2 s to ensure that each dataset was created from the same set of EEGs (Fig **4**A). All segments were zero-padded to match the original input length of 2.0 s. The standard 3D CNN maintained *>*95% accuracy down to 0.5-s segments but declined to 86% accuracy with 0.1-s segments (Fig **4**B). Notably, BiCL pre-training enabled the 3D CNN to sustain *>*90% accuracy down to 0.5-s segments in both fine-tuning scenarios, achieving 94% under low-data and 91% under class imbalance. These results demonstrate that seizure and non-seizure states can be effectively distinguished at sub-second durations, even with limited seizure data. Since high accuracy was maintained with 0.5-s segments, we used the 0.5-s parietal cortex dataset for the remaining results in this work.

**Figure 4:**
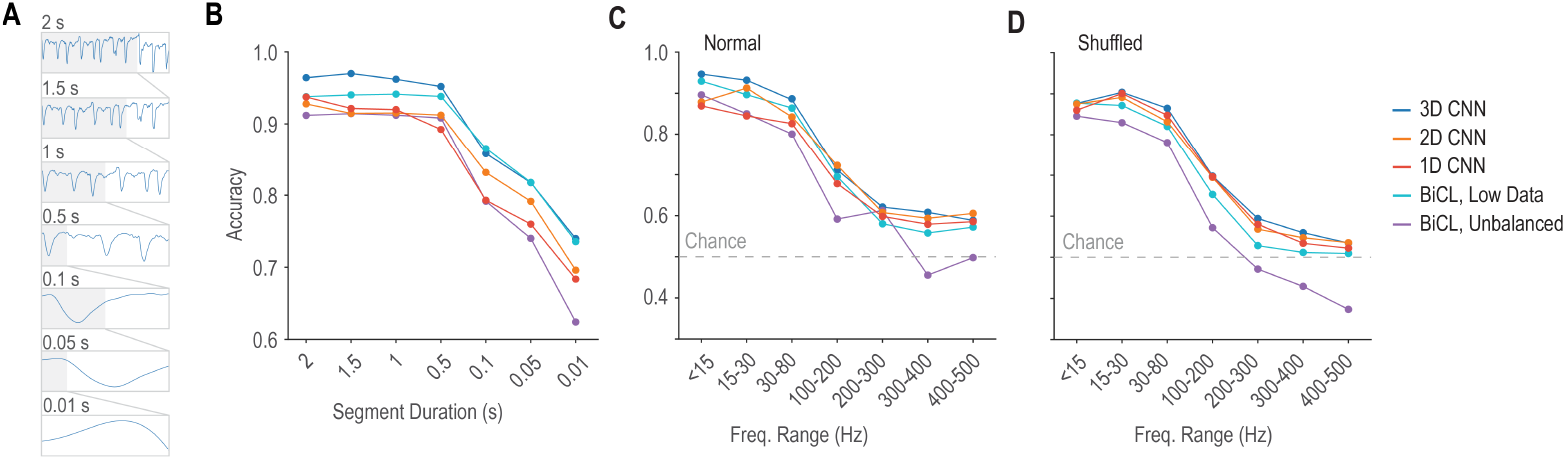
Performance with different segment lengths and at different frequency bands. (A) Example seizure trace (blue) exemplifying the effect of using different segment lengths: 2.0, 1.5, 1.0, 0.5, 0.1, 0.05, and 0.01 s. The length of each segment appears in the top left corner and the grayed area corresponds to the length of the next segment from top to bottom. (B) Mean accuracy of the models at different segment duration. Zero-padding was applied to all the segments to maintain the original input length. (C) Mean accuracy of the models when the EEG data is band-pass filtered across the ranges and the temporal structure is preserved. (D) Mean accuracy of the models when the data is first band-pass filtered and the samples in the signal are randomly shuffled. Color legend is the same for B, C, and D.

The EEG signatures of seizures can appear across a broad frequency spectrum, ranging from as low as 0.5 Hz to above 70 Hz.^48^ To investigate the frequency components that the CNNs depend upon, the data was band-pass filtered using the following ranges: *<*15 Hz, 15–30 Hz, 30–80 Hz, 100–200 Hz, 200–300 Hz, 300–400 Hz, and 400–500 Hz. The 3D, 2D, and 1D CNNs were then trained and tested on each filtered dataset. For the BiCL 3D CNN, each dataset was used for fine-tuning and testing under the two scenarios previously described. The results showed that all architectures performed best within the frequency bands less than 80 Hz (Fig **4**C). Specifically, in the *<*15 Hz band, the 3D CNN achieved 95% accuracy, while the BiCL 3D CNN attained 93% and 90% accuracy under low-data and class-imbalanced scenarios, respectively. Performance declined substantially at higher frequencies; in the 100–200 Hz band, accuracies decreased to 71%, 70%, and 59% for the same models. These findings demonstrate that CNNs predominantly rely on low-frequency information for seizure detection.

We next examined whether seizure detection depends on temporal patterns or merely on the statistical properties of the EEG data. After dividing each band-pass filtered signal into segments, we randomly shuffled each segment to determine if the presence of a seizure is embedded in the statistics of the data, such as the mean and variance, or the temporal organization.^49^ Across all frequency bands, models trained on shuffled data achieved accuracies that were comparable to—or only slightly lower than—those of the original data (Fig **4**D). The only exception was the 1D CNN, which exhibited small performance gains in the 15–30 Hz, 30–80 Hz, and 100–200 Hz bands. These results suggest that temporal patterns play a minor role in seizure detection. Moreover, since both shuffled and un-shuffled models achieved their highest accuracies in the ranges less than 80 Hz, all further experiments in this work are conducted using EEG data low-pass filtered at 80 Hz, a range that aligns well with what is reliably available in commercial EEG systems.^50^

The robustness of seizure detection algorithms to signal degradation is critical for real-world clinical applications, where EEG patterns that characterize a seizure can be conflated with artifacts.^51^ We evaluated our approach under two perturbation scenarios: noise and downsampling. To first test the performance of our approach under conditions with low signal-to-noise ratios, we perturbed the data by adding zero-mean Gaussian noise to each EEG recording, where the noise standard deviation was scaled proportionally to the variability of the signal. Specifically, we defined *σ*_noise_ = *k × σ*_signal_, where *k* ∈ {0, 1, 2, 5, 10, 20, 50, 100} represents the noise scaling factor.^52^ This approach ensured that noise amplitude scaled appropriately with signal characteristics while preserving the original signal mean. Separate 1D, 2D, and standard 3D CNNs were trained and tested for each noise level. For the BiCL 3D CNN, we performed fine-tuning under both low-data and class-imbalance. As expected, all architectures exhibited performance degradation with increasing noise (Fig **5**A). The standard 3D CNN maintained an accuracy *>*90% up to the moderate noise level of *k* = 2. Moreover, the 1D and 2D CNNs showed steeper decreases in accuracy, dropping to *<*80% at *k* = 100 while the 3D CNN exhibited accuracies *>*85% at all noise levels.

**Figure 5:**
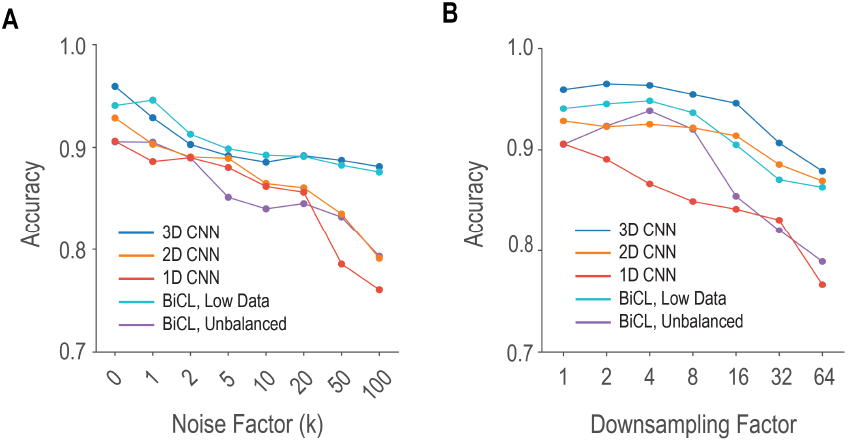
Performance when adding noise or downsampling. (A) Mean accuracy with noise added, where the noise has mean 0 and a standard deviation equal to the standard deviation of each EEG, scaled by factors of 0, 1, 2, 5, 10, 20, 50, and 100. (B) Mean accuracy after downsampling with factors of 1, 2, 4, 8, 16, 32, and 64.

Since EEG signals are sparse, downsampling is often used to reduce computational complexity while preserving the essential characteristics of the original signal.^53^ Downsampling disproportionately affects seizure segments, which contain steep voltage deflections that require adequate temporal resolution. We evaluated the effect of temporal resolution on seizure detection by applying downsampling factors of 1, 2, 4, 8, 16, 32, and 64 to the original 2713 Hz recordings. To maintain consistent input dimensions across all conditions, we zero-padded down-sampled segments to preserve the original length. Independent models were trained and evaluated at each downsampling level. The standard 3D CNN achieved an accuracy of *>*90% up to a factor of 32, suggesting that fine temporal details play a minimal role in model predictions when sufficient training data is available (Fig **5**B). However, the BiCL 3D CNN with low-data fine-tuning resulted in an accuracy of 87% at a factor of 32, indicating greater reliance on temporal resolution when training samples are limited. For the class-imbalance scenario, the highest accuracy of 94% was observed at a moderate downsampling factor of 4. In general, CNNs are often touted as requiring minimal pre-processing and have been used to directly classify raw EEG signals.^54^ However, in the absence of ample training data, smoothing the input (via downsampling) leads to more prominent seizure features relative to noise, making the limited seizure samples more salient and improving minority-class detection.

Rather than overfitting to the idiosyncratic features of a single mouse, an ideal model should learn common electrophysiological patterns. Given our dataset of 10 mice, we use a leave-one-subject-out cross-validation approach, whereby each model was trained on data from nine mice and subsequently tested on the remaining mouse. This procedure was repeated until each of the 10 mice served as the testing set 10 times, with the average accuracy over all trials computed per model for each mouse (Fig **6**B). For the BiCL 3D CNN, we simulated two clinically relevant scenarios. The low-data scenario used a random 10% of each of the nine training mice’s data, while the class-imbalanced scenario used 10% of seizure segments and 90% of non-seizure segments, randomly sampled from each training mouse. For both scenarios, the testing data was the complete dataset of the tenth mouse. Across all mice, the standard 3D CNN demonstrated superior cross-subject generalization, achieving 93% mean accuracy with low variability (*σ* = 6%). Notably, mice M01, M02, and M03 had the smallest datasets, therefore prediction errors can have a large effect on performance metrics (Fig **6**A). In contrast, mouse M10 had the longest total duration of seizure data, and removing this dataset results in a training set with the shortest total duration of seizure data, which may explain why these four mice showed the lowest accuracies in our evaluation. Moreover, the BiCL 3D CNN maintained robust performance, attaining average accuracies of 90% and 91% for the low-data and class-imbalanced scenarios, respectively. These results indicate that our approach reliably identifies seizures in subjects on which it was not trained, thereby capturing fundamental neural dynamics rather than subject-specific anomalies, even with sub-second temporal windows.

**Figure 6:**
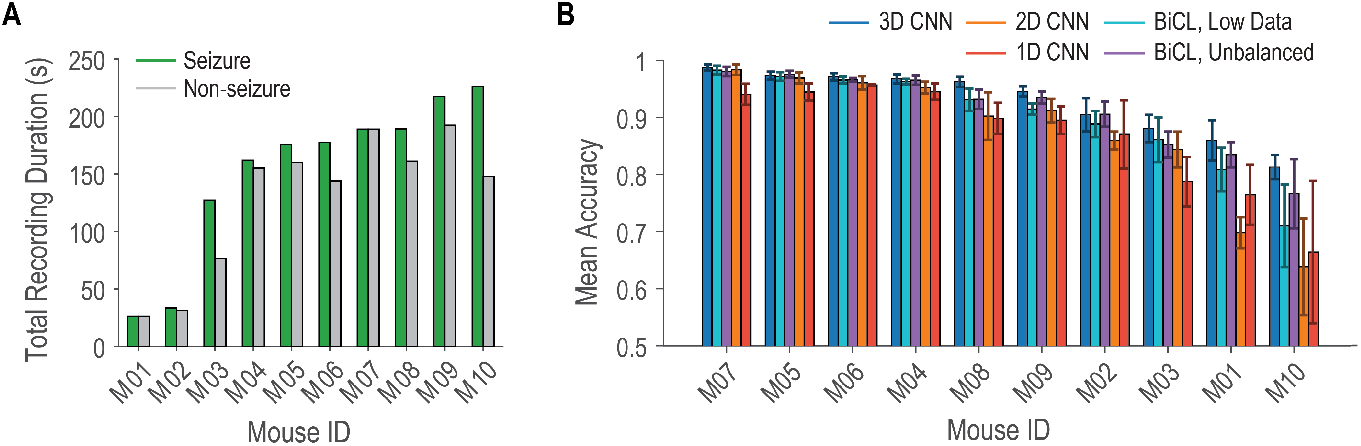
Performance in a leave-one-subject-out condition. (A) Total recording duration for seizures and non-seizures for the parietal EEG segments across all mice. (B) Mean accuracy for each mouse over 10 trials of leave-one-subject-out cross-validation.

## Discussion

We have shown that, on single-channel parietal cortex data, the standard 3D CNN achieved the highest accuracy, outperforming both the 2D CNN, which applies the CWT as a pre-processing step, and the 1D CNN, which operates directly on raw signals. This result shows that incorporating the CWT directly within the network architecture allows for more robust feature extraction. In general, the primary cause of delayed epilepsy diagnosis is a failure to recognize seizures by patients, families, and healthcare providers.^55^ The low false negative rate is significant in clinical settings, where failing to detect a seizure event could lead to inadequate treatment, especially in emergency settings where seizures are frequently under-detected and under-reported.^56^ Interestingly, the 3D CNN showed only a minimal performance increase when data from both cortices was used compared to data from just the parietal cortex. Such robustness indicates that seizure detection can be conducted using one channel, which is relevant as widespread use of multichannel EEG is impeded by cost and setup and interpretation times.^57^

In many real-world settings, it is difficult to obtain large amounts of labeled data. While there are multiple publicly available EEG seizure datasets, the sparse occurrence of seizures (occurring in *<*1% of data in many datasets), as well as factors like seizure duration, electrode placement, and the number of channels, make seizure identification using machine learning difficult.^58^ The use of contrastive pre-training addresses these challenges by learning generalizable representations from unlabeled data. The superior performance of BiCL over CPC in single-channel, low-data scenarios suggests that leveraging both past and future contexts enables the model to capture more comprehensive seizure patterns. Moreover, contrastive learning successfully extracted transferable seizure representations despite the substantial domain shift between pre-training and finetuning datasets: human vs. mouse, scalp vs. epidural recording, electrode configurations, and referencing schemas. This cross-domain generalization demonstrates that our approach captures fundamental signatures of seizure activity, enabling robust detection even with minimal task-specific training data or severe class imbalance.

The investigation into temporal resolution revealed insights about the minimum duration needed for reliable seizure detection. The ability of the 3D CNN to maintain accuracy exceeding 95% with sub-second segments suggests that the network can identify seizure characteristics even from brief snippets of EEG data. Although our study focused on retrospective analysis rather than realtime implementation, our results suggest applicability to online detection systems that could immediately notify family members, caregivers, and emergency units, reducing seizure-related injuries and improving quality of life.^59^ Furthermore, when the EEGs were band-pass filtered, all architectures performed best in the *<*80 Hz range, which aligns with clinical insights that epileptic seizures give rise to changes in the delta, theta, alpha, and beta frequency bands.^60^ The sharp decline in performance at frequencies above 100 Hz suggests that high-frequency oscillations may not provide consistent discriminative features across a diverse seizure dataset. This finding has practical implications for system design since focusing on lower frequency bands could reduce sampling rate requirements without sacrificing accuracy. Moreover, the disruption of temporal order through shuffling showed that temporal organization plays a less important role in seizure detection, indicating that the statistical properties of the EEG signal are the primary drivers of model performance.

Additionally, our investigation into the impact of noise and downsampling revealed several practical advantages of the proposed approach. The resilience of the 3D CNN to the addition of Gaussian noise indicates practical viability in clinical environments, where EEG recordings are susceptible to artifact contamination.^61^ Moreover, the retention of high accuracy under moderate downsampling further suggests that a focus on broader patterns is sufficient for seizure detection. This result reduces computational demands and facilitates the deployment of such models in resource-constrained systems. In epilepsy diagnosis, EEG data may differ significantly across subjects and clinical centers, and the performance of a classifier trained on data from multiple subjects generally declines when applied to new subjects.^62^ Our cross-subject evaluation demonstrated that the 3D CNN generalizes effectively to unseen subjects, capturing universal neural patterns rather than overfitting to subject-specific features. This generalizability suggests that our approach can be applied to new subjects without requiring retraining or calibration.

We have shown that incorporating the CWT as a trainable layer is an effective and robust approach for feature extraction. Future work will focus on evaluating our method under more challenging conditions to explore its clinical viability. We will assess performance across distinct seizure types with heterogeneous EEG signatures, including focal and generalized seizures. We also plan to validate the framework using data acquired in realistic clinical settings, including recordings where quality may be compromised by electrical interference, patient movement, and other sources of noise. Additionally, we will extend our evaluation to longer, continuous recordings that include sleep-wake transitions, as seizure patterns can vary significantly between different states of consciousness.^63^ Ultimately, the strong performance and generalization capabilities demonstrated in this work indicate that integrating wavelet-based feature extraction with deep learning offers a promising pathway towards reliable, automated seizure detection in clinical practice and research settings.

## Acknowledgments

This work was generously supported by the JPB Foundation; the Picower Institute for Learning and Memory; George J. Elbaum (MIT ‘59, SM ‘63, PhD ‘67), Mimi Jensen, Diane B. Greene (MIT, SM ‘78), Mendel Rosenblum, Bill Swanson, annual donors to the MIT/MGH BASCIC Fund; and the NIH Awards P01 GM118269 and R01 NS123120 (to E.N.B.). We are also grateful to Mark Olchanyi for helpful feedback on the manuscript.

